# Contrast-induced changes in chemical exchange saturation transfer MRI differentiate tumor progression from pseudoprogression

**DOI:** 10.64898/2026.05.01.722099

**Authors:** Blake Benyard, Narayan Datt Soni, Anshuman Swain, Nishi Srivastava, Sunil Kumar Khokhar, Akila Ramesh, Junyoung Shin, Dushyant Kumar, Ravi Prakash Reddy Nanga, Nadir Yehya, Yi Fan, Ravinder Reddy, Mohammed Haris

## Abstract

Tumor pseudo-progression (PsP) refers to an initial increase in tumor size or the appearance of new lesions. These pseudo-progressive lesions are predominantly composed of infiltrative inflammatory cells, such as macrophages. This phenomenon commonly occurs in patients undergoing radiation therapy or immunotherapy and typically indicates a positive treatment response. However, it often leads to premature treatment cessation due to misinterpretation as disease progression. Non-invasive imaging biomarkers capable of distinguishing pseudo-progression from true progression would greatly aid in treatment decision-making. In our preliminary study, we explored the potential of gadoterate meglumine (Gd-DOTA, a macrocyclic Gd-contrast) in combination with amine chemical-exchange saturation transfer (amine-CEST) imaging to differentiate tumor from radiation necrosis by assessing Gd-DOTA uptake by infiltrating immune cells, such as macrophages. To evaluate whether amine-CEST, in combination with Gd-DOTA, can differentiate macrophages from cancer cells, we incubated them with Gd-DOTA for 30 minutes. Subsequently, the cells were processed, and amine-CEST imaging was performed on a 9.4 Tesla preclinical scanner. Upon treatment with Gd-DOTA, we did not observe a significant change in amine-CEST contrast in F98 cells compared with untreated cells, whereas treated macrophages exhibited a marked decrease (∼40%) in amine-CEST signal compared with untreated macrophages. This reduction in signal was attributed to the uptake of Gd-DOTA by macrophages, which notably shortened water T_1_ relaxation, thereby quenching the amine-CEST signal. Conversely, cancer cells showed no appreciable change in the amine-CEST signal, indicating no Gd-DOTA uptake. Furthermore, to validate that T_1_ shortening influences amine-CEST signal, cancer cells were also treated with manganese chloride (MnCl_2_) for 30 minutes. The uptake of MnCl_2_ by cancer cells similarly induced T_1_ shortening, as observed in macrophages, resulting in a decrease in the amine-CEST signal from these cells. Next, we performed the amin-CEST imaging on F98 tumor-bearing rats and radiation necrotic rats. Post-injection with Gd-DOTA showed no appreciable change in the amine-CEST contrast in the tumor-bearing rat, whereas a significant decrease in contrast was observed in the radiation necrotic rat. This further demonstrates that no change in the amine-CEST contrast in tumor-bearing rats is due to cancer cells failing to take up Gd-DOTA. The decrease in amine-CEST contrast in radiation-treated rats reflects the uptake of Gd-DOTA by macrophages infiltrating the radiation-necrotic regions. This straightforward imaging approach holds promise for clinical translation. It offers a novel method for characterizing pseudo-progressive lesions and monitoring diverse treatment responses in cancer patients using standard clinical scanners.

## Introduction

Tumor treatments, such as immunotherapy, chemotherapy, and radiation therapy (RT), can induce inflammation within the tumor regions, which can be misinterpreted as true tumor progression (TP) (*1-13*). Specifically, TP is characterized by growing tumor regions with new tumor cells, whereas pseudoprogression (PsP) mimics this growth due to an extracellular environment infiltrated with immune cells, such as macrophages, that respond to RT or other treatments. About 50% of patients exhibit either TP or PsP (*14*). Differentiating TP from PsP is crucial in oncology, especially for brain tumors and cancers treated with radiation therapy (RT), immunotherapy, or targeted therapies ^6-19^. TP indicates that the cancer is growing or worsening, which may necessitate changes in treatment, such as switching to a different therapy, intensifying current treatment, or exploring clinical trials. Conversely, PsP reflects a temporary increase in tumor size or imaging changes that can mimic progression but are indicative of a treatment response^1-9^. Misinterpreting PsP as TP could lead to unnecessary changes in the treatment plan, potentially halting a beneficial therapy. Accurate differentiation between these two conditions using a non-invasive imaging approach is crucial for making informed treatment decisions. This differentiation also helps manage patient anxiety, prevent harmful interventions, accurately evaluate treatment responses, maintain the integrity of clinical trials, and inform long-term prognosis and follow-up plans.

Current imaging modalities, such as positron emission tomography (PET) and conventional MRI, including contrast enhancement, have several drawbacks when used to evaluate immunotherapeutic responses and distinguish between PsP and TP (*14*). PET is limited by additional scanning requirements, lower spatial resolution, and a lack of a reference standard for histologic and radiologic confirmation, which can delay accurate PsP diagnosis and affect treatment scheduling. Furthermore, varying tracer use across studies complicates comparisons between experiments.

Treatments induce local tissue inflammation leading to edema and increased vascular permeability (*14*). This results in fluid accumulation, which is evident on T_2_-weighted magnetic resonance imaging (MRI) sequences and appears as a new lesion. Immunotherapeutic responses have also been monitored using diffusion-weighted imaging (DWI) and its quantitative measure, the apparent diffusion coefficient (ADC), with some limitations, as ADC values can be inaccurate due to differences in cell density between necrotic and solid tumor regions (*15-17*). Gadolinium-based contrast agents (GBCAs) and manganese (Mn_2_^+^) have been used in MRI to enhance contrast by reducing T_1_ relaxation times (*18*). Specifically, GBCAs are being widely used in clinics to image tumors in human patients. However, conventional T_1_-weighted imaging with GBCA often highlights the entire region, making it difficult to distinguish TP from PsP (*19*). Other techniques, such as contrast clearance analysis, assess GBCA clearance in tumor lesions to differentiate PsP from TP, but the lengthy acquisition process can lead to patient discomfort and motion artifacts, and may introduce errors in the interpretation of findings (*20*).

Due to these limitations, there is a clear unmet need for advanced, non-invasive imaging techniques with high-resolution capabilities that can reliably differentiate between TP and PsP. Emerging imaging methods, such as chemical exchange saturation transfer (CEST), have the potential to detect early metabolic alterations in tumors and to characterize TP and PsP (*21-39*). Amine-CEST is a technique for indirectly detecting mobile metabolites and macromolecules, such as proteins, that have exchangeable amine protons. This technique has been used in both clinical patients and preclinical animal models, demonstrating increased amine-CEST contrast in brain tumors compared with normal brain tissue. However, because both macrophages and tumor cells exhibit CEST effects, distinguishing between them with standard CEST methods is challenging. In this study, we used amine-CEST MRI in combination with GBCA to determine whether we could distinguish macrophages, indicative of PsP, from tumor cells, which represent TP. In addition, we have applied these methods to distinguish tumors from radiation necrosis in rats *in vivo*.

## Materials and Methods

Amine-CEST imaging was performed on bovine serum albumin (BSA), F98 cancer cells, and macrophages in the presence and absence of gadoterate meglumine (Gd-DOTA). Also, amine-CEST maps were acquired from cancer cells incubated with manganese chloride (MnCl_2_). BSA phantom MRI experiments were performed using a 3T whole body scanner (MAGNETOM Prisma, Siemens Healthcare, Erlangen, Germany) with a body transmit/32-channel receive proton phased-array head coil, as well as a 7T whole body scanner (MAGNETOM Terra, Siemens Healthcare, Erlangen, Germany) with a Nova Medical volume coil transmit/32-channel receive proton phased-array head coil, at 25 °C. All cell line and animal experiments were performed using a 20 mm diameter ^1^H transceiver volume head coil (m2m Imaging) on a 9.4 T horizontal magnet interfaced with an Advance III HD console (Bruker BioSpin, Germany) at 25 °C. Animal study was performed under an approved Institutional Animal Care and Use Committee (IACUC) protocol.

### Bovine serum albumin preparation and amine-CEST MRI

A 10% (weight/volume) BSA solution was prepared in phosphate-buffered saline (PBS, pH = 7.0). We prepared two solution phantoms in 10 mm NMR tubes - one doped with 0.1mM of Gd-DOTA and another without Gd-DOTA. The amine-CEST imaging was performed on both 3T and 7T human scanners. The amine-CEST acquisition parameters for 3T were: field-of-view (FOV) = 100 × 100 mm^2^, matrix size = 128 × 128, slice thickness = 5 mm, TE = 4 ms, averages = 2, shot TR = 8 s, saturation pulse of B_1,rms_ = 2 µT, and saturation duration = 1s.

At 7T, the amine-CEST acquisition parameters were: FOV = 100 × 100 mm^2^, matrix size = 128 × 128, slice thickness = 5 mm, TE = 1.79 ms, averages = 2, shotTR = 8 s, saturation pulse of B_1,rms_ = 2 µT, and saturation duration = 1s.

The CEST images were acquired at evenly spaced saturation offset frequencies from -5 to +5 ppm (relative to the water resonance) with a step-size of 0.2 ppm. An image without pre-saturation was acquired as a reference for the CEST asymmetry calculation.

### Cancer cell preparation

F98 cancer cells were cultured in Roswell Park Memorial Institute (RPMI) 1640 medium supplemented with 10% fetal bovine serum and incubated at 37°C in a 5% CO_2_ environment. The cultures were maintained by splitting every three days. For the experiment, we prepared three groups of cells, each containing approximately 50 × 10^6^ cells. One group served as the control, while the other two groups were incubated with 20 µL of Gd-DOTA and MnCl_2_, respectively. After a 30-minute incubation, the cells were washed 3 times with PBS to remove residual contrast agent, then centrifuged to obtain cell pellets for imaging.

### Macrophages preparation

Human-derived monocytes were cultured in RPMI 1640 medium supplemented with 2 mM L-glutamine, 10% fetal bovine serum (FBS), and 1% antibiotic-antimycotic (AA) solution, along with 50 ng/mL macrophage colony-stimulating factor and 25 ng/mL interleukin-10 (IL-10). The cells were incubated at 37°C in a 5% CO_2_ environment. The medium was changed every two days until the monocytes differentiated into inflammatory macrophages. Once 95% of the monocytes had differentiated into macrophages, the cells were collected and divided into two groups. One group was treated with Gd-DOTA, followed by a 30-minute incubation period, while the other group remained untreated as a control. The cells were collected and washed with PBS three times to remove residual Gd-DOTA and centrifuged to obtain the pallet for scanning at 9.4T animal scanner.

### Brain tumor model

Five Sprague-Dawley rats (∼6 weeks old, ∼140 g) were used to generate a tumor-bearing glioma model. Briefly, rats were anesthetized with isoflurane, and their scalp hair was removed using a hair trimmer and hair removal cream. Rats were positioned with their heads fixed on the stereotaxic apparatus equipped for a continuous supply of isoflurane to induce a surgical plane of anesthesia. Bupivacaine (under the scalp; 2 mg/kg) and meloxicam (subcutaneous; 2 mg/kg) were administered as analgesics. After sterilizing the scalp, an incision was made to expose the skull. Following stereotaxic navigation to 3 mm lateral (right) and 3 mm posterior to bregma, a burr hole was drilled to inject a 5 μl suspension containing ∼50,000 gliosarcoma (F98) cells in phosphate buffer into the cerebral cortex (2 mm deep) using a Hamilton syringe and a 32-gauge needle. These rats underwent MR imaging 3 weeks after tumor cell implantation.

### Radiation necrosis model

Rats were irradiated using a Small Animal Radiation Research Platform (SARRP; Xstrahl, Suwanee, GA) delivering photon X-ray irradiation at a nominal dose rate of 3.80 Gy/min. Animals received 40 Gy per fraction, administered once per week over a three-week treatment course (total dose: 120 Gy). Prior to irradiation, rats were anesthetized with isoflurane and immobilized using a customized bed to ensure reproducible positioning. Beam geometry and targeting were planned using onboard cone-beam CT to localize the region of interest and minimize exposure to surrounding structures. Dose delivery and field uniformity were verified using the vendor-supplied treatment planning system and in-house dosimetry protocols. Following the irradiation regimen, animals were monitored longitudinally for delayed radiation effects. Necrotic tissue formation was observed approximately nine months after completion of the final fraction.

### 9.4T MR Imaging

Both cell line and rat brain imaging were performed on a 9.4T horizontal bore preclinical MR scanner (Bruker) and a 30 mm-diameter proton-volume coil (m2m Imaging Corp., OH). Animals were kept under anesthesia (1.5% isoflurane in 1 liter/min oxygen), and their body temperature was maintained by circulating air generated and blown through a heater (SA Instruments, Inc., Stony Brook, NY). Respiration and body temperature were continuously monitored using an MRI-compatible small animal monitoring system (SA Instruments, Inc., Stony Brook, NY).

At first, a localizer was acquired for sample positioning, followed by a T_1_-weighted FLASH (TE = 4 ms, TR = 498 ms, four averages, 16 slices) and T2-weighted RARE (TE1/TE2 = 33/121 ms, TR = 3s, two averages, 16 slices, rare factor = 6. CEST imaging on cell lines was performed using the following parameters: in-plane resolution = 0.156 mm x 0.156 mm, field-of-view = 20 × 20 mm^2^, matrix size = 128 × 128, slice thickness = 1mm, gradient-echo readout TR = 3 ms, TE = 4 ms, averages = 2, T1 delay = 6 s, saturation pulse of B_1,rms_ = 4 µT, and saturation duration = 1s. For animal imaging, the same parameters were used, except that B_1rms_ = 1 µT and saturation duration = 3s. An image without pre-saturation was acquired as a reference for the CEST asymmetry calculation. The CEST images were acquired at evenly spaced saturation offset frequencies from - 5 to +5 ppm (relative to the water resonance) with a step-size of 0.2 ppm. To correct for B_0_ field inhomogeneity, B_0_ maps were acquired using water saturation shift referencing (WASSR), with images collected from 0 to ±1 ppm (step-size = 0.1 ppm, saturation pulse of B_1,rms_ = 0.1 µT, and 200-ms duration)^51^. After baseline imaging, an intravenous injection (IV) of a gadolinium-based contrast agent (Gadoterate meglumine, Dotarem, France; 0.1 mmol/kg) (Gd-DOTA) was administered through a pre-inserted catheter in the tail vein. To monitor changes in signal intensity, T_1_-weighted images and CEST images were acquired again on the same slice 30 minutes post-injection.

### Data post-processing

Data processing was performed using in-house MATLAB (version R2019a) scripts, as described previously by Benyard et al.^52^. The amine-CEST asymmetry ratio was computed from CEST weighted images at a frequency offset of (Δω = +2.0 ppm) using the following equation:

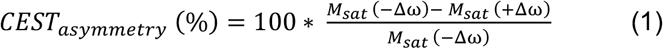

where M_Sat_ (± Δω) is the equilibrium magnetization achieved with a train of saturation pulses applied at ±2 ppm relative to the water peak.

## Results

Figure 1 provides a conceptual framework for understanding the differential interactions between contrast agents, macrophages, and cancer cells. It is hypothesized that macrophages possess a highly efficient mechanism for the uptake of Gd-DOTA, a commonly used MRI contrast agent. This capability may be attributed to macrophages’ innate properties and their ability to internalize various substances. In contrast, cancer cells exhibit significantly lower Gd-DOTA uptake. This discrepancy might stem from the differences in cellular metabolism, membrane permeability, or receptor expression between macrophages and cancer cells. The differential uptake can have important implications for developing imaging methods to image inflammatory responses in cancers and other diseases.

**Figure 1.**
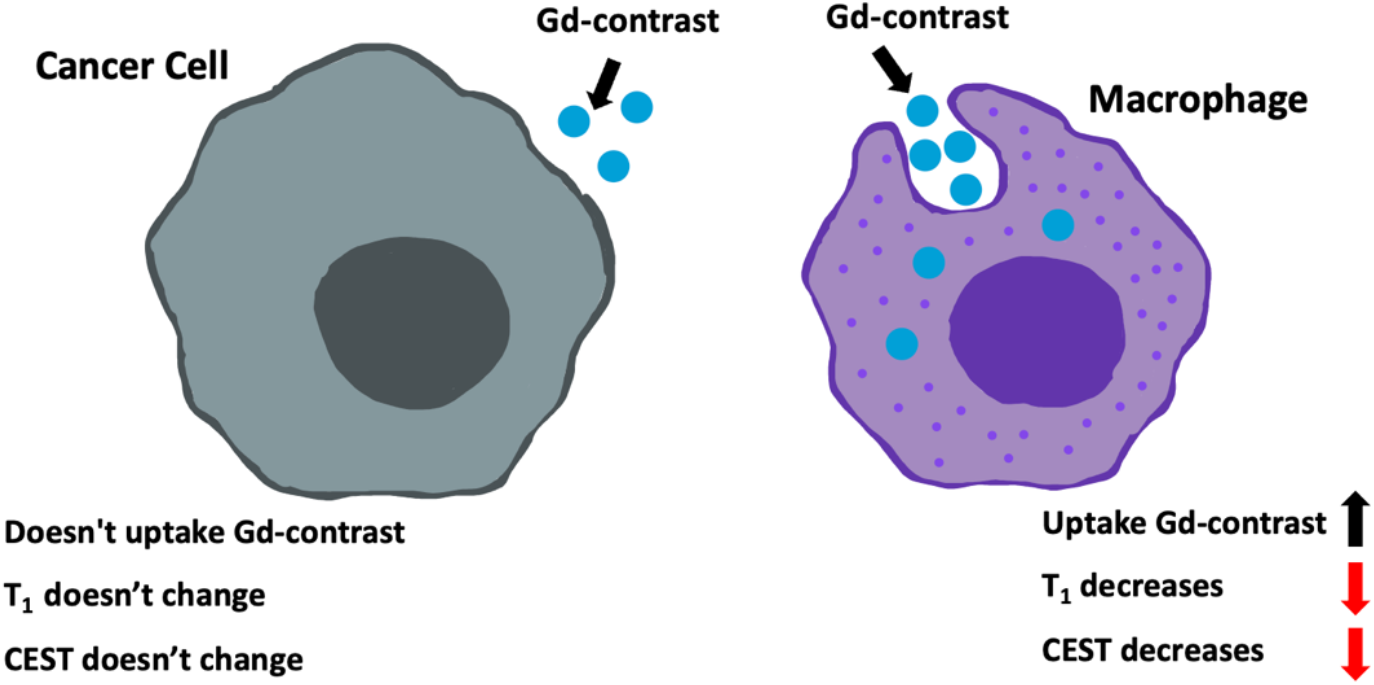
The mechanism describing the theory behind the experiment. Here it is illustrated that the cancer cells, representing the intracellular space, do not uptake the contrast agent (gadolinium). Macrophages, which occupy the extracellular space, take up gadolinium, thereby reducing T_1_ and CEST contrast.

For proof of concept, we performed amine-CEST experiments on BSA phantoms (10% BSA) with and without Gd-DOTA. Representative Z-spectra at 7T (**Fig. 2a**) demonstrate a significant reduction in amine-CEST contrast at 2 ppm in the presence of Gd-DOTA. This reduction is characterized by a loss in asymmetry and a narrowing of the Z-spectral width compared to the BSA phantom without Gd-DOTA. The amine-CEST maps of BSA phantoms at 3T and 7T show decreased amine-CEST contrast in the Gd-DOTA-doped samples (**Fig. 2b & c**). Specifically, at 3T, the undoped BSA sample exhibited approximately 30% contrast, while the Gd-DOTA-doped sample showed no detectable amine-CEST signal. Similarly, at 7T, the undoped BSA solution displayed around 35% amine-CEST contrast, whereas the Gd-DOTA-doped solution exhibited less than 5% contrast.

**Figure 2.**
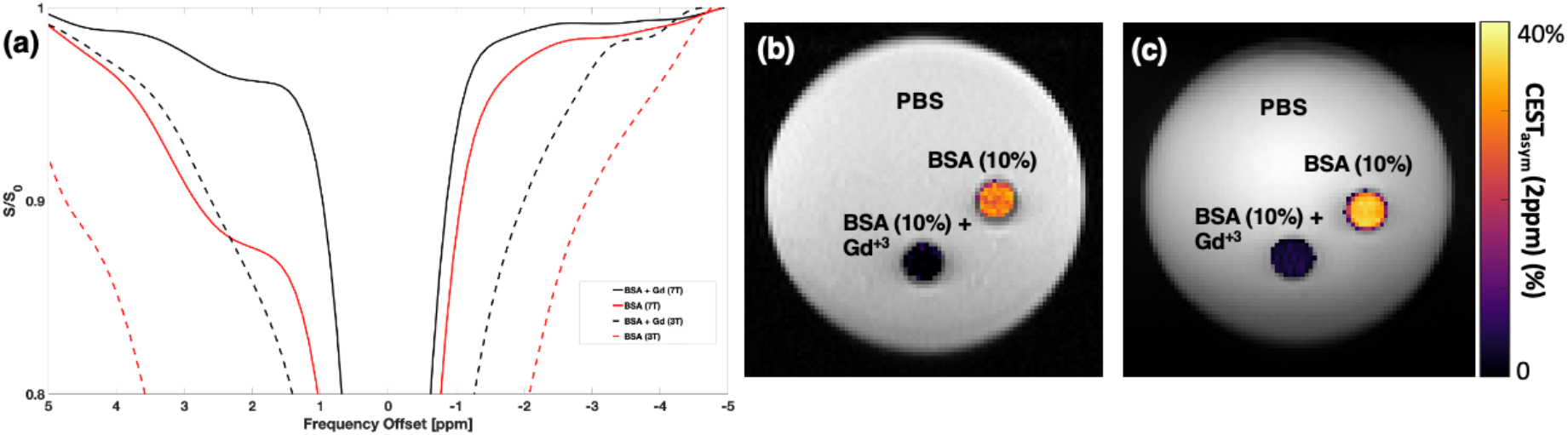
Z-spectrum from the BSA phantom at 3T (dotted lines) and 7T (a). This Z-spectrum shows that the sample with Gd-DOTA does not exhibit CEST contrast at 2 ppm. Amine-CEST maps of the BSA phantoms at 3T (b) and 7T (c). CEST maps show a clear drop in contrast when combined with the Gd-DOTA contrast agent, both at 3T and 7T.

Figure 3a displays the amine-CEST maps of F98 cancer cells incubated with and without Gd-DOTA, showing no appreciable change in amine-CEST contrast (∼33%) between the two groups. The corresponding asymmetry plots from 0 to 5 ppm (**Fig. 3b**) further confirm the lack of significant contrast alteration, indicating that F98 cancer cells do not uptake Gd-DOTA within 30 minutes of incubation.

**Figure 3.**
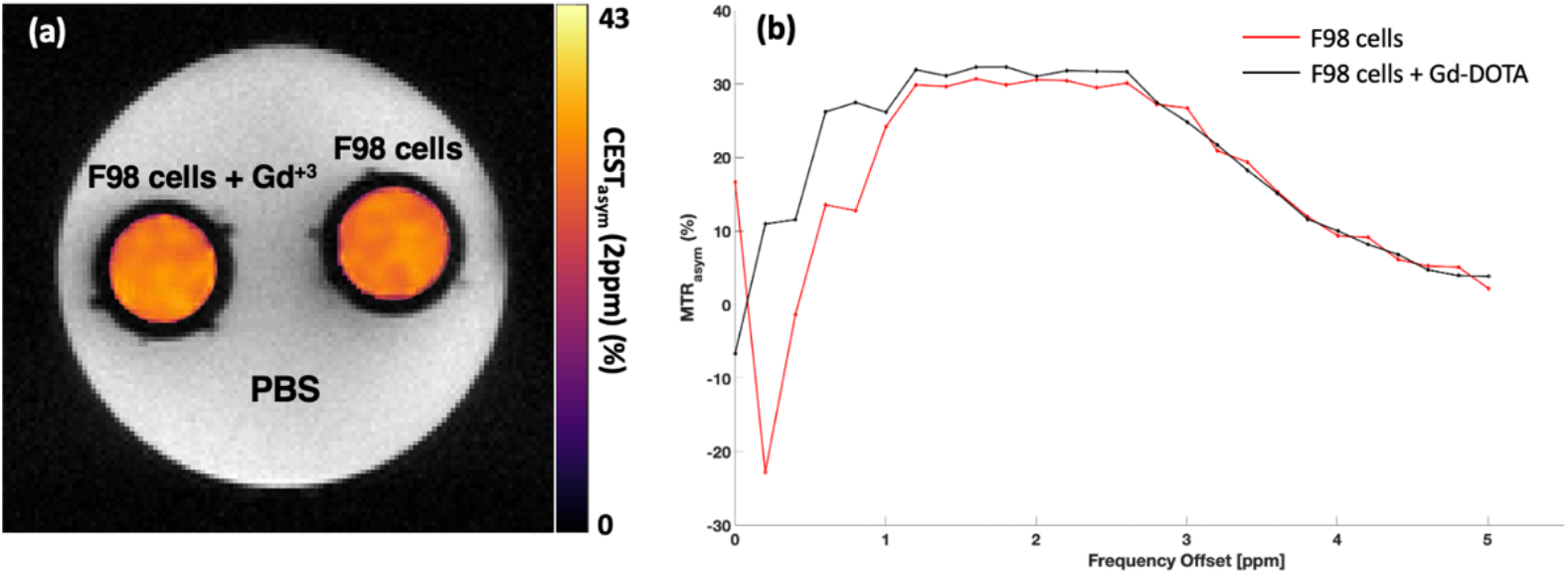
Amine-CEST maps from the F98 cancer cells incubated with and without Gd-DOTA (a) and the corresponding MTR asymmetry plots generated from ROI’s placed in the cells (b). The CEST maps show no change in amine-CEST contrast between the two sets of cells in phantoms with and without Gd-DOTA. The asymmetry plots also show no appreciable difference in asymmetry at 2 ppm.

Figure 4a shows amine-CEST maps from macrophages with and without Gd-DOTA. Macrophages incubated with Gd-DOTA demonstrated a marked reduction in amine-CEST contrast (∼30%) compared to those without Gd-DOTA (∼70%). The faster decline in MTR asymmetry for Gd-DOTA-doped macrophages is depicted in Figure 4b, highlighting the impact of Gd-DOTA on macrophage amine-CEST contrast.

**Figure 4.**
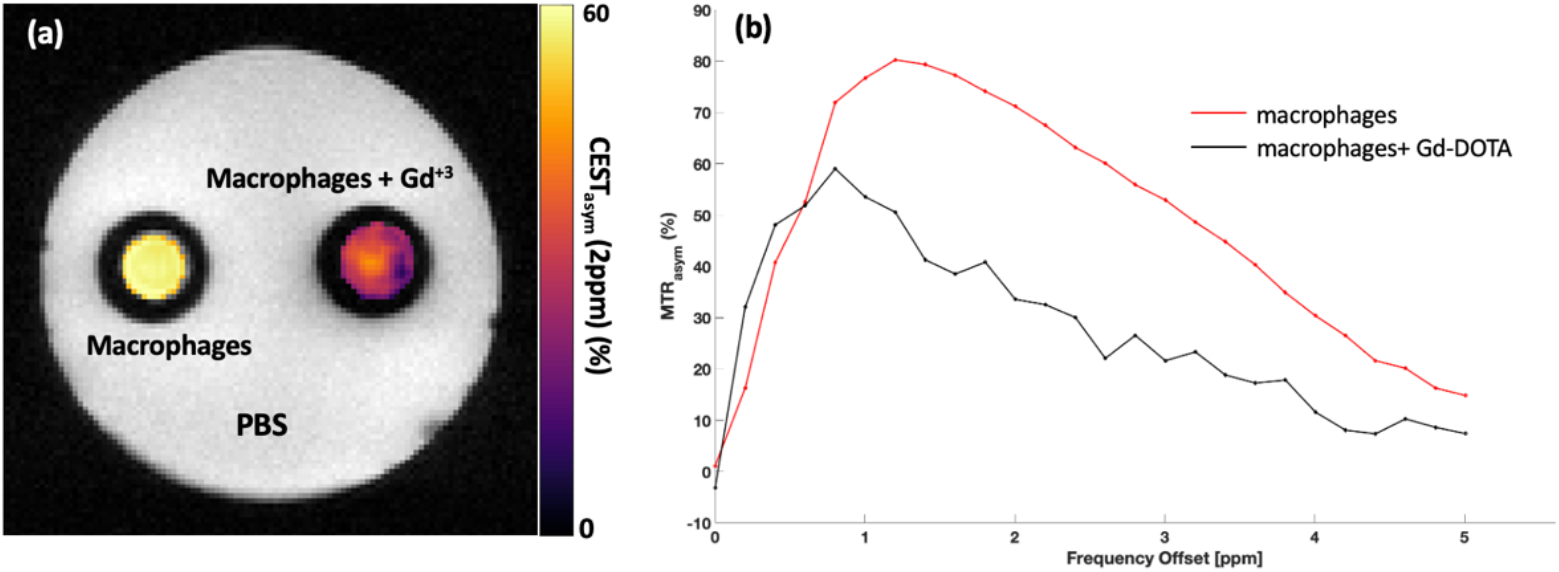
The amine-CEST maps from the macrophage samples with and without Gd-DOTA (a) and the corresponding MTR asymmetry plots generated from ROI’s placed in the cells (b). These maps also show that the macrophages doped with Gd-DOTA uptook the agent and consequently have significantly less CEST contrast (∼50%). The asymmetry plots further illustrate that there is a significant difference in asymmetry across the frequency offsets.

We further tested if amine-CEST contrast in cancer cells could be quenched by incubating F98 cells with MnCl2. As shown in Figure 5a, the MnCl2-treated F98 cells exhibited a drastic reduction in amine-CEST contrast (∼1%) compared to untreated cells (∼33%). The corresponding MTR asymmetry plots in Figure 5b reveal that MnCl_2_ not only quenches the amine-CEST signal at 2 ppm but also disrupts the signal across the entire spectrum.

**Figure 5.**
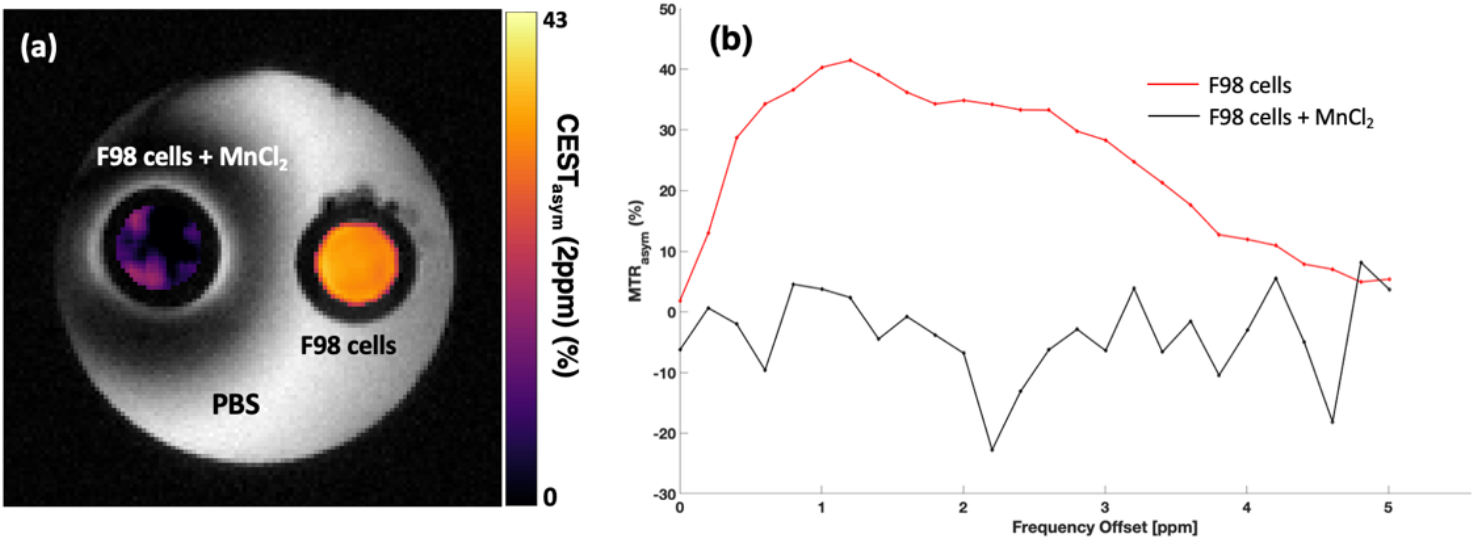
Amine-CEST maps from the F98 cancer cells incubated with and without MnCl_2_ (a) and the corresponding MTR asymmetry plots generated from ROI’s placed in the cells (b). The CEST maps show a significant drop (∼40%) in contrast compared with cells incubated with MnCl_2_. The asymmetry plots further illustrate that there is a significant difference in asymmetry across the frequency offsets.

Figure 6 shows the amine-CEST maps from the animal models of glioma (**Fig. 6a**) and radiation necrosis (**Fig. 6b**) pre- and post-Gd-DOTA infusion. After the infusion of Gd-DOTA, there was no difference in amine-CEST contrast from the tumorous tissue; however, a significant decrease in amine-CEST contrast was observed in the radiation-induced necrotic brain region (**Fig. 6c**).

**Figure 6.**
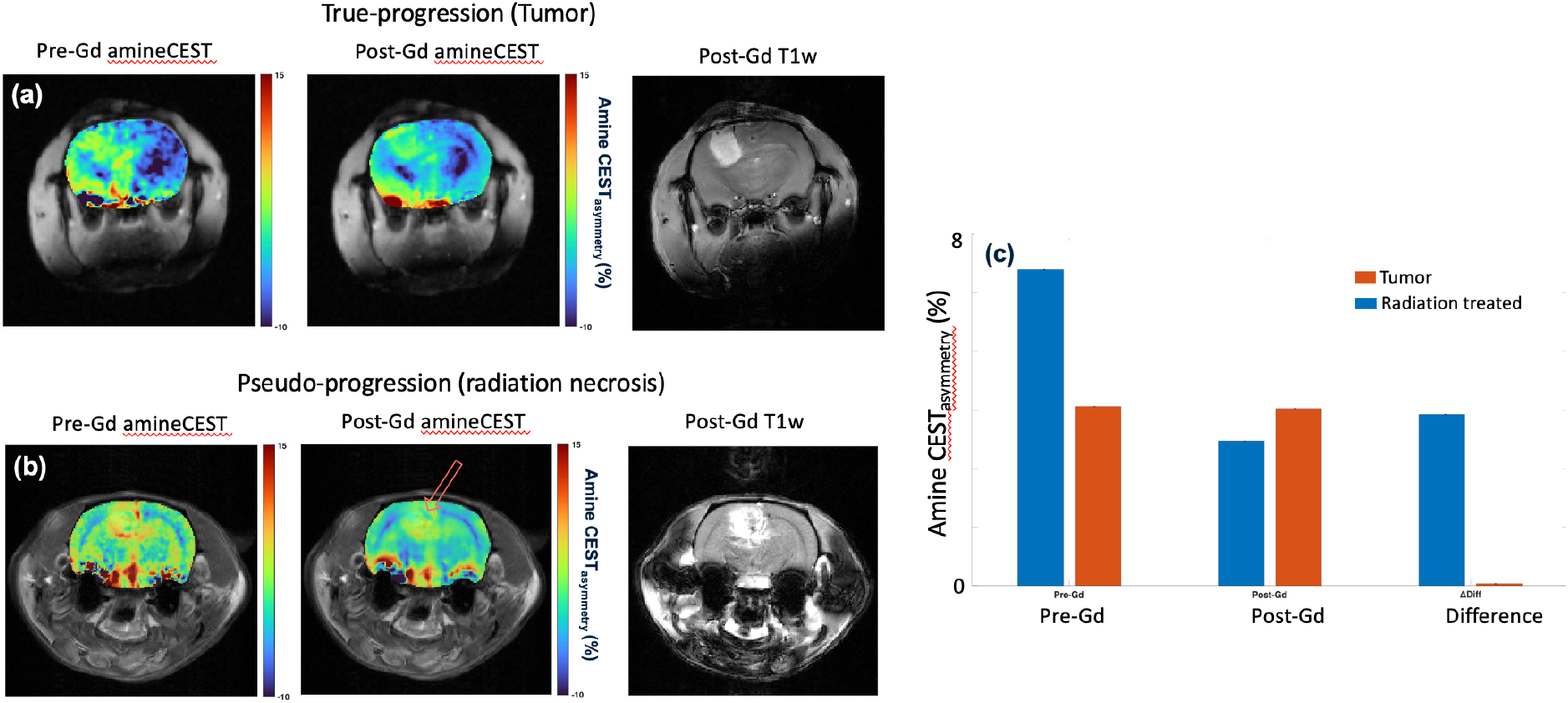
In vivo amine-CEST maps in a rat glioma model (a) and a radiation necrosis model pre- and post-Gd administration (b). The bar plot shows the change in amineCEST contrast post-Gd-DOTA infusion in both tumor and necrotic regions (c).

## Discussion

To the best of our knowledge, this study represents the first application of amine-CEST MRI to successfully differentiate between cancer cells and macrophages. Prior literature has established that gadolinium-based contrast agents (GBCAs) are not taken up in the intracellular compartments of cancer cells but are readily absorbed by extracellular compartments. Our findings are consistent with these observations, as we demonstrate that amine-CEST MRI depends on T_1_ relaxation times using both BSA phantoms and macrophages. The BSA phantoms and macrophages doped with GBCAs exhibited a significant reduction in amine-CEST contrast due to T_1_ shortening, underscoring the influence of Gd-DOTA on MRI signals. This reduction in CEST contrast confirms that T_1_ shortening, induced by GBCA, directly affects the CEST signal.

Furthermore, we validated the effect of T_1_ shortening on amine-CEST in cancer cells incubated with MnCl_2_. The observed significant decrease in amine-CEST contrast in the presence of MnCl_2_ suggests that manganese ions effectively quench the CEST signal, likely due to active transport through calcium channels into cancer cells^28^. This mechanism, in which MnCl_2_ induces T_1_ shortening, has been previously documented in studies involving prostate cancer cells at 7T and human embryonic stem cells at 3T. These findings further support the utility of GBCAs for distinguishing different cell types *in vitro*.

We observed no change in the amine-CEST signal in the rat glioma model after Gd-DOTA infusion, indicating that the tumor cells did not take up the Gd contrast agent (**supplementary figure 1**). In contrast, there was a significant decrease in amine-CEST signal in the radiation-induced necrotic regions, suggesting the presence of infiltrating immune cells, including macrophages. These cells efficiently took up the gadolinium contrast, resulting in the observed decrease in amine-CEST contrast. The findings of this study also align with previous research using APT CEST imaging, as shown by Torrealdea et al., who demonstrated that the contrast from tumor tissue pre- and post-Gd administration (**supplementary figure 2**) remained similar^55^. This lack of contrast change further confirms that lesions composed of cancer cells do not uptake GBCAs, thereby resulting in no alteration in CEST contrast. Similarly, Tee et al. reported minimal changes in APT CEST contrast in glioma patients pre- and post-administration of a chelated Gd contrast agent, reinforcing the observation that cancer cells do not absorb GBCAs^56^. However, their study does not confirm which form of chelated Gd they used.

Studies have demonstrated the uptake of linear forms of gadolinium contrast agents in both normal and cancerous cells. In the current study, however, we used a macrocyclic gadolinium agent (Gd-DOTA), which has been shown to be more stable and is not taken up by either normal or cancer cells. In another study, Multiple sclerosis (MS) patients who received macrocyclic Gd-agents showed increased T_1_ hyperintensity in the globus pallidus and dentate nucleus (*40*). It was suggested that this could be identified as an active inflammation in the brain of MS patients (*40*).

Given that cancer cells do not take up GBCAs, their T_1_ relaxation times remain unaffected, resulting in similar amine-CEST signals regardless of the presence of Gd-DOTA. However, when cancer cells were incubated with MnCl_2_, a significant decrease in CEST contrast was observed, consistent with the T_1_-shortening effect of manganese ions, which are actively taken up through calcium channels. This observation opens new avenues for selectively targeting cancer cells and immune cell infiltration with T_1_-modifying agents to enhance the specificity of MRI-based diagnostics.

The approach presented in this study holds promise for differentiating TP from PsP in clinical settings and for monitoring macrophage immune responses. Given that macrophages and other inflammatory cells are major contributors to PsP lesions and readily uptake foreign substances, differential uptake of Gd-DOTA by macrophages versus cancer cells could serve as a distinguishing feature between TP and PsP lesions. Our results suggest that amine-CEST contrast would remain unchanged in TP lesions, while a significant decrease in amine-CEST contrast could be expected in PsP lesions post-GBCA administration.

Although this study focused on the feasibility of using exogenous contrast agents to differentiate TP from PsP in animal models, the principles demonstrated here can be translated to human studies, laying the groundwork for future investigations. Additionally, while we utilized amine-CEST MRI, the findings are applicable to other endogenous CEST methods, including APT CEST. The broader applicability of these results could enhance the utility of CEST MRI in various oncological imaging scenarios.

## Conclusion

In summary, our study demonstrates that cancer cells do not take up the macrocyclic form of Gd-contrast (Gd-DOTA), whereas macrophages do, leading to differential effects on amine-CEST contrast. This differentiation has significant implications for improving MRI accuracy in distinguishing TP from PsP, potentially leading to better-informed treatment decisions and improved outcomes in clinical oncology.

## Figures

**Supplementary Figure 1.**
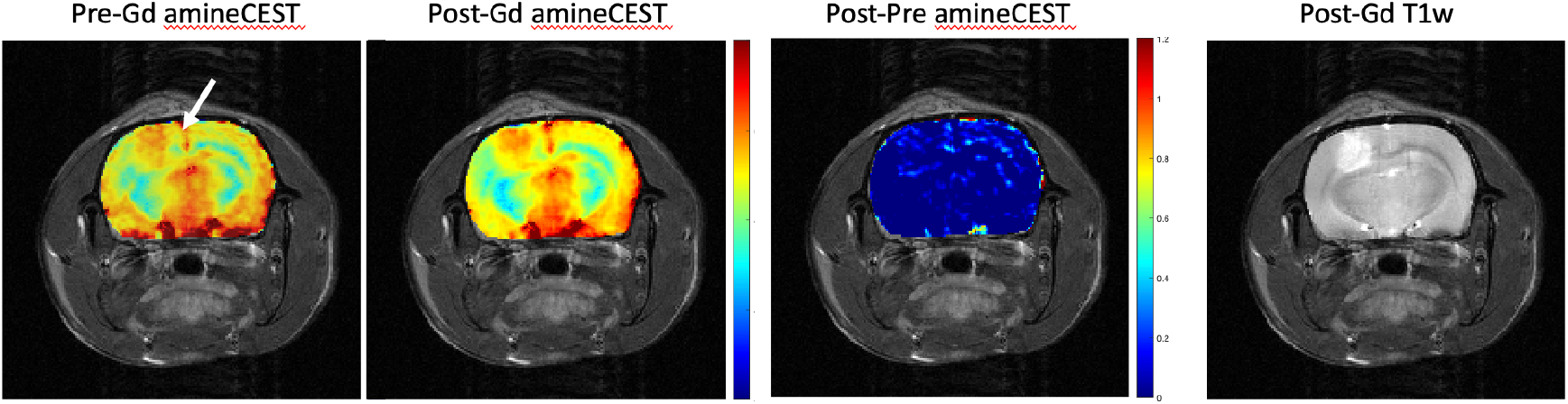
AmineCEST contrast maps from a rat glioma model before and after the infusion of Gd-DOTA. The difference image clearly shows that there was no change in the amineCEST contrast in either the tumor or the normal-appearing brain regions. The post-contrast T1-weighted image reveals the tumor as a hyperintense area.

**Supplementary Figure 2.**
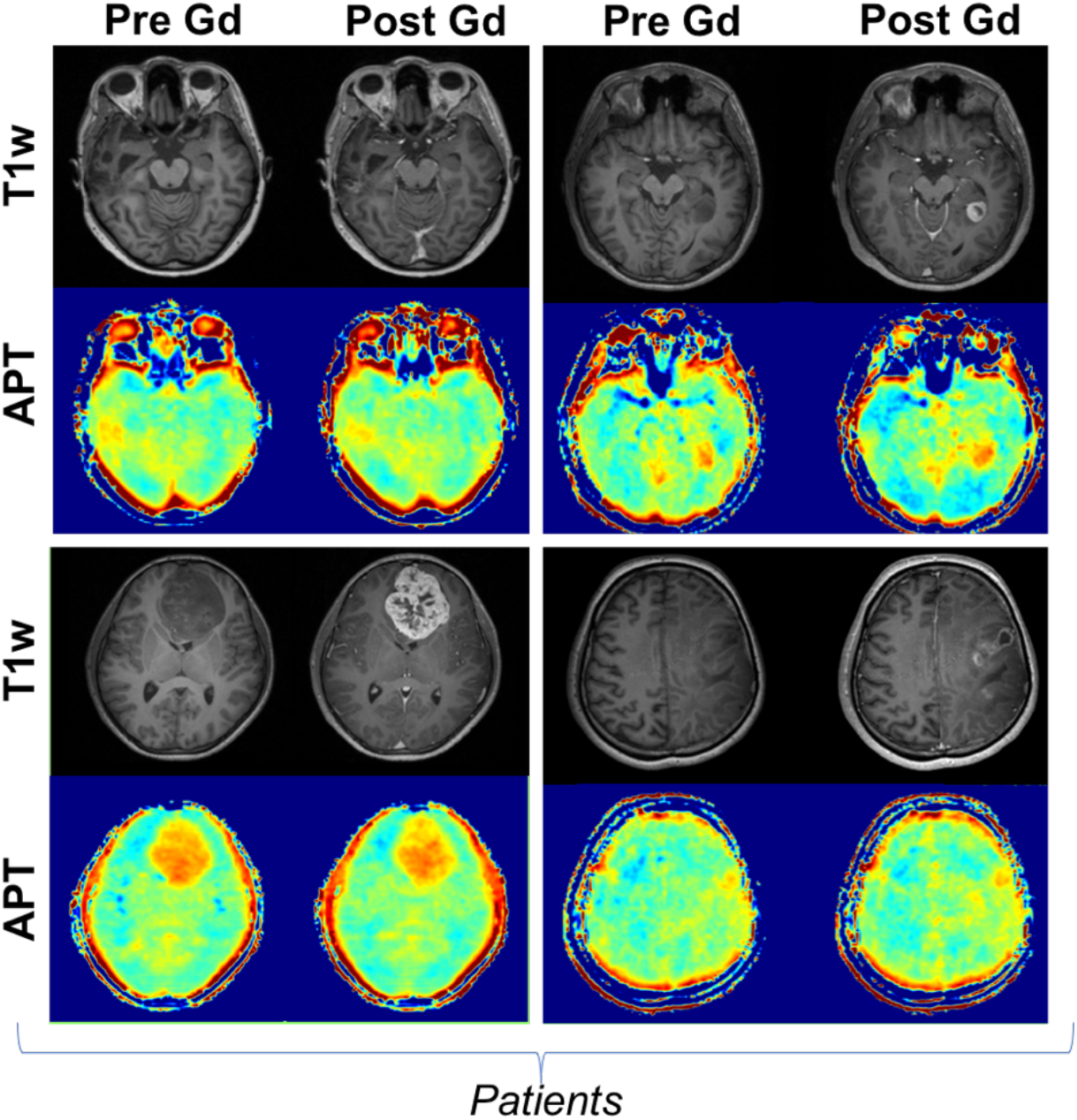
Pre- and post-Gd APT-CEST and T1w images of four patients show no change in the APT contrast from both the tumor and normal appearing brain regions. In all patients, tumor appears as a hyperintense area on post-contrast T1-weighted images. Adapted from Torrealdea, Francisco & Hearle, J & Evans, et al. APT-CEST post Gadolinium. Should it be avoided? Comparison of pre- & post-Gadolinium CEST on glioma at 3T. Intl. Soc. Mag. Reson. Med. 26 (2018).

## Acknowledgements

Research reported in this publication was supported by the National Institutes of Health under award numbers R01AG091760, R01AG063869, and RF1AG087306 and by the National Institutes of Health’s National Institute of Biomedical Imaging and Bioengineering under award number P41EB029460. The authors thank Emily Cento, Zhilin Chen, Max A. Eldabbas, and Emileigh Maddox of the Human Immunology Core and the Division of Transfusion Medicine and Therapeutic Pathology at the Perelman School of Medicine at the University of Pennsylvania for providing de-identified, human-derived monocytes that were purified from healthy donor apheresis using StemCell RosetteSep™ kits. The HIC is supported in part by NIH P30 AI045008 and P30 CA016520.

## Notes

### Competing Interest Statement

The authors have declared no competing interest.

### Summary of Updates

We updated the author list based on the contribution

## References

1. P. D. Delgado-Lopez, E. Rinones-Mena, E. M. Corrales-Garcia, Treatment-related changes in glioblastoma: a review on the controversies in response assessment criteria and the concepts of true progression, pseudoprogression, pseudoresponse and radionecrosis. Clin Transl Oncol 20, 939–953 (2018).

2. M. R. Gilbert et al., Dose-dense temozolomide for newly diagnosed glioblastoma: a randomized phase III clinical trial. J Clin Oncol 31, 4085–4091 (2013).

3. D. Hanahan, R. A. Weinberg, Hallmarks of cancer: the next generation. Cell 144, 646–674 (2011).

4. A. Malmstrom et al., Temozolomide versus standard 6-week radiotherapy versus hypofractionated radiotherapy in patients older than 60 years with glioblastoma: the Nordic randomised, phase 3 trial. Lancet Oncol 13, 916–926 (2012).

5. W. Roa et al., Abbreviated course of radiation therapy in older patients with glioblastoma multiforme: a prospective randomized clinical trial. J Clin Oncol 22, 1583–1588 (2004).

6. W. Roa et al., International Atomic Energy Agency Randomized Phase III Study of Radiation Therapy in Elderly and/or Frail Patients With Newly Diagnosed Glioblastoma Multiforme. J Clin Oncol 33, 4145–4150 (2015).

7. M. Sneeggen, N. A. Guadagno, C. Progida, Intracellular Transport in Cancer Metabolic Reprogramming. Front Cell Dev Biol 8, 597608 (2020).

8. R. Stupp et al., Effects of radiotherapy with concomitant and adjuvant temozolomide versus radiotherapy alone on survival in glioblastoma in a randomised phase III study: 5-year analysis of the EORTC-NCIC trial. Lancet Oncol 10, 459–466 (2009).

9. J. J. Vredenburgh et al., Bevacizumab plus irinotecan in recurrent glioblastoma multiforme. J Clin Oncol 25, 4722–4729 (2007).

10. W. Wick et al., Temozolomide chemotherapy alone versus radiotherapy alone for malignant astrocytoma in the elderly: the NOA-08 randomised, phase 3 trial. Lancet Oncol 13, 707–715 (2012).

11. T. A. Wynn, A. Chawla, J. W. Pollard, Macrophage biology in development, homeostasis and disease. Nature 496, 445–455 (2013).

12. O. L. Chinot et al., Bevacizumab plus radiotherapy-temozolomide for newly diagnosed glioblastoma. N Engl J Med 370, 709–722 (2014).

13. S. M. Davidson, M. G. Vander Heiden, Critical Functions of the Lysosome in Cancer Biology. Annu Rev Pharmacol Toxicol 57, 481–507 (2017).

14. S. C. Thust, M. J. van den Bent, M. Smits, Pseudoprogression of brain tumors. J Magn Reson Imaging 48, 571–589 (2018).

15. H. H. Chu et al., Differentiation of true progression from pseudoprogression in glioblastoma treated with radiation therapy and concomitant temozolomide: comparison study of standard and high-b-value diffusion-weighted imaging. Radiology 269, 831–840 (2013).

16. Y. Li et al., Advanced Imaging Techniques for Differentiating Pseudoprogression and Tumor Recurrence After Immunotherapy for Glioblastoma. Front Immunol 12, 790674 (2021).

17. Y. S. Song et al., True progression versus pseudoprogression in the treatment of glioblastomas: a comparison study of normalized cerebral blood volume and apparent diffusion coefficient by histogram analysis. Korean J Radiol 14, 662–672 (2013).

18. C. A. Massaad, R. G. Pautler, Manganese-enhanced magnetic resonance imaging (MEMRI). Methods Mol Biol 711, 145–174 (2011).

19. J. S. Young, N. Al-Adli, K. Scotford, S. Cha, M. S. Berger, Pseudoprogression versus true progression in glioblastoma: what neurosurgeons need to know. J Neurosurg 139, 748–759 (2023).

20. R. Bodensohn et al., MRI-based contrast clearance analysis shows high differentiation accuracy between radiation-induced reactions and progressive disease after cranial radiotherapy. ESMO Open 7, 100424 (2022).

21. H. Y. Heo et al., Whole-brain amide proton transfer (APT) and nuclear overhauser enhancement (NOE) imaging in glioma patients using low-power steady-state pulsed chemical exchange saturation transfer (CEST) imaging at 7T. J Magn Reson Imaging 44, 41–50 (2016).

22. T. Jin, P. Wang, X. Zong, S. G. Kim, MR imaging of the amide-proton transfer effect and the pH-insensitive nuclear overhauser effect at 9.4 T. Magn Reson Med 69, 760–770 (2013).

23. C. K. Jones et al., Nuclear Overhauser enhancement (NOE) imaging in the human brain at 7T. Neuroimage 77, 114–124 (2013).

24. G. Liu, X. Song, K. W. Chan, M. T. McMahon, Nuts and bolts of chemical exchange saturation transfer MRI. NMR Biomed 26, 810–828 (2013).

25. H. Mehrabian, K. L. Desmond, H. Soliman, A. Sahgal, G. J. Stanisz, Differentiation between Radiation Necrosis and Tumor Progression Using Chemical Exchange Saturation Transfer. Clin Cancer Res 23, 3667–3675 (2017).

26. H. Mehrabian, S. Myrehaug, H. Soliman, A. Sahgal, G. J. Stanisz, Evaluation of Glioblastoma Response to Therapy With Chemical Exchange Saturation Transfer. Int J Radiat Oncol Biol Phys 101, 713–723 (2018).

27. J. E. Meissner et al., Early response assessment of glioma patients to definitive chemoradiotherapy using chemical exchange saturation transfer imaging at 7 T. J Magn Reson Imaging 50, 1268–1277 (2019).

28. D. Paech et al., Nuclear Overhauser Enhancement imaging of glioblastoma at 7 Tesla: region specific correlation with apparent diffusion coefficient and histology. PLoS One 10, e0121220 (2015).

29. D. Paech et al., Nuclear overhauser enhancement mediated chemical exchange saturation transfer imaging at 7 Tesla in glioblastoma patients. PLoS One 9, e104181 (2014).

30. S. Regnery et al., Chemical exchange saturation transfer MRI serves as predictor of early progression in glioblastoma patients. Oncotarget 9, 28772–28783 (2018).

31. Y. Shen et al., Imaging of nuclear Overhauser enhancement at 7 and 3 T. NMR Biomed 30, (2017).

32. X. Tang et al., Nuclear Overhauser Enhancement-Mediated Magnetization Transfer Imaging in Glioma with Different Progression at 7 T. ACS Chem Neurosci 8, 60–66 (2017).

33. Y. K. Tee, M. J. Donahue, G. W. Harston, S. J. Payne, M. A. Chappell, Quantification of amide proton transfer effect pre- and post-gadolinium contrast agent administration. J Magn Reson Imaging 40, 832–838 (2014).

34. P. C. van Zijl, N. N. Yadav, Chemical exchange saturation transfer (CEST): what is in a name and what isn’t? Magn Reson Med 65, 927–948 (2011).

35. J. Windschuh et al., Correction of B1-inhomogeneities for relaxation-compensated CEST imaging at 7 T. NMR Biomed 28, 529–537 (2015).

36. Y. Wu et al., 3D APT and NOE CEST-MRI of healthy volunteers and patients with non-enhancing glioma at 3 T. MAGMA 35, 63–73 (2022).

37. M. Zaiss et al., Relaxation-compensated CEST-MRI of the human brain at 7T: Unbiased insight into NOE and amide signal changes in human glioblastoma. Neuroimage 112, 180–188 (2015).

38. X. Y. Zhang et al., MR imaging of a novel NOE-mediated magnetization transfer with water in rat brain at 9.4 T. Magn Reson Med 78, 588–597 (2017).

39. J. Zhou et al., Differentiation between glioma and radiation necrosis using molecular magnetic resonance imaging of endogenous proteins and peptides. Nat Med 17, 130–134 (2011).

40. F. G. Moser et al., High Signal Intensity in the Dentate Nucleus and Globus Pallidus on Unenhanced T1-Weighted MR Images: Comparison between Gadobutrol and Linear Gadolinium-Based Contrast Agents. AJNR Am J Neuroradiol 39, 421–426 (2018).

